# Vegetation formation in *Staphylococcus aureus* endocarditis inversely correlates with *RNAIII* and *sarA* expression in invasive clonal complex 5 (CC5) isolates

**DOI:** 10.1101/2022.04.20.488955

**Authors:** Kyle J. Kinney, Jessica M. Stach, Katarina Kulhankova, Matthew Brown, Wilmara Salgado-Pabón

## Abstract

Infective endocarditis (IE) is one of the most feared and lethal diseases caused by *Staphylococcus aureus.* Once established, the infection is fast-progressing and tissue destructive. *S. aureus* of the clonal complex 5 (CC5) commonly cause IE yet are severely understudied. IE results from bacterial colonization and formation of tissue biofilms (known as vegetations) on injured or inflamed cardiac endothelium. *S. aureus* IE is promoted by adhesins, coagulases, and superantigens, with the exotoxins and exoenzymes likely contributing to tissue destruction and dissemination. Expression of the large repertoire of virulence factors required for IE and sequelae is controlled by complex regulatory networks. We investigated the temporal expression of the global regulators *agr* (*RNAIII*), *rot*, *sarS*, *sarA*, *sigB*, and *mgrA* in 8 invasive CC5 isolates and established intrinsic expression patterns associated with IE outcomes. We show that vegetation formation, as tested in the rabbit model of IE, inversely correlates with *RNAIII* and *sarA* expression during growth in Todd-Hewitt broth (TH). Large vegetations with severe sequelae arise from strains with high-level expression of colonization factors but slower transition towards expression of the exotoxins. Overall, strains proficient in vegetation formation, a hallmark of IE, exhibit lower expression of *RNAIII* and *sarA*. Simultaneous high expression of *RNAIII, sarA*, *sigB*, and *mgrA* is the one phenotype assessed in this study that fails to promote IE. Thus, *RNAIII* and *sarA* expression that provides for rheostat control of colonization and virulence genes, rather than an on and off switch, promote both vegetation formation and lethal sepsis.

## Introduction

*Staphylococcus aureus* is an opportunistic pathogen that colonizes the mucosal surfaces and skin of 30-40% of the human population (Spaulding et al., 2013). It causes a wide range of illnesses, from very common superficial skin and soft tissue infections to life-threatening, invasive and fulminant diseases such as sepsis and infective endocarditis (IE) (Lowy, 1998; Boucher et al., 2010; Bergin et al., 2017). Currently, *S. aureus* IE develops in 5-32% of patients with *S. aureus* bacteremia and overall accounts for the majority of IE cases in high-income countries (Bergin et al., 2017; Fernández-Hidalgo et al., 2018; Moreau et al., 2018). *S. aureus* IE is also increasingly common in settings where viridans streptococci are the primary etiologic agent of IE due to a high burden of congenital heart disease (Bergin et al., 2017). Left-sided, native valve IE is the most common manifestation of *S. aureus* endocarditis (Werdan et al., 2014). Once established, *S. aureus* IE is fast-progressing and tissue destructive, leading to metastatic infections and severe systemic complications that result in 22-66% lethality even with modern medical and surgical interventions (Bergin et al., 2017; Fernández-Hidalgo et al., 2018). Without treatment, *S. aureus* IE is 100% lethal (Lerche et al., 2021). In fact, *S. aureus* infection represents an independent risk factor associated with IE in-hospital mortality (Fowler et al., 2005; Fernández-Hidalgo et al., 2012; Selton-Suty et al., 2012).

*S. aureus* IE results from bacterial infection of injured or inflamed cardiac endothelium, predominantly the heart valves (Werdan et al., 2014). *S. aureus* adhesion and colonization triggers endothelial inflammation, expression of tissue factors and fibronectin, and host factor aggregation (e.g. fibrin, fibrinogen, platelets), promoting development and growth of vegetations pathognomonic of IE. For valve colonization, *S. aureus* binds to extracellular matrix proteins via surface adhesins anchored to the cell wall collectively known as MSCRAMMs (microbial surface components recognizing adhesive matrix molecules) and secreted adhesins known as SERAMs (secretable expanded repertoire adhesive molecules) (Patti et al., 1994; Bergin et al., 2017). *S. aureus* adhesins that contribute to cardiac valve colonization and vegetation formation include ClfA (clumping factor or fibrinogen-binding protein A) and FnBPA (fibronectin-binding protein A) (McDevitt et al., 1995; Sinha et al., 1999; Kerrigan et al., 2008). Vegetation growth and maturation is promoted by adhesins, coagulases, and superantigens. The adhesins and coagulases trigger platelet activation and aggregation. These include ClfA and FnBPs but also protein A, SdrE (serine–aspartate repeat-containing protein E), IsdB (iron-regulated surface determinant protein B), Eap (extracellular adherence protein), and coagulases Coa and vWbp (von-Willebrand factor binding protein) (Que et al., 2005; Heying et al., 2007; Bertling et al., 2012; Vanassche et al., 2012). The superantigens of the *egc* (enterotoxin gene cluster), TSST-1 (toxic shock syndrome toxin-1), and SEC (staphylococcal enterotoxin C) and the sphingomyelin *β*-toxin alter endothelial cell function and promote vegetation growth (Stach et al., 2016; Kulhankova et al., 2018; Kinney et al., 2022; Tran et al., 2022). Superantigens, hemolysins, and exoenzymes are thought to contribute to the aggressive and tissue destructive nature of *S. aureus* IE (King et al., 2016; Tran et al., 2022).

*S. aureus* utilizes intricate and complex regulatory networks to control the expression of the large repertoire of cell-surface and secreted virulence factors that promote host colonization, immune evasion, persistence, and disease development (Jenul and Horswill, 2019). This network includes multiple global regulators of virulence gene expression with some of the most prominent being the accessory gene regulator (*agr*) quorum-sensing system, the transcriptional regulators SarA, SarS, Rot, and MgrA, and the alternative sigma factor SigB (Recsei et al., 1986; Somerville and Proctor, 2009). The cell-density dependent control of adhesins and colonization factors versus exotoxins and spreading factors promotes the transition from a colonization to a dissemination phenotype (Fechter et al., 2014). Regarding *S. aureus* IE, strain-to-strain variation in gene regulation is likely one factor contributing to differences in disease presentation.

The most common *S. aureus* clonal groups isolated from IE patients include clonal complex (CC)5, CC8, CC30, and CC45 (Xiong et al., 2009; Nienaber et al., 2011). The CC5 background is of particular interest because of its overall association with persistent bacteremia and hematogenous complications, its high frequency in infections around the globe, and its association with deregulation of the Agr system (Xiong et al., 2009; Fernández-Hidalgo et al., 2018). Further interest in *S. aureus* CC5 isolates stems from their significant cause of health care-associated, methicillin-resistant infections in the Western world and from being the principal genetic background associated with full vancomycin resistance (Challagundla et al., 2018). Yet, most of the understanding of *S. aureus* pathogenesis and global regulation of virulence gene expression comes from the study of USA300 strains, which commonly activate the Agr system at high levels (Li et al., 2009; Montgomery et al., 2010; Grundstad et al., 2019). Recent studies have emphasized the need to acquire a greater understanding of the molecular and pathogenic mechanisms characteristic of this clonal group (King et al., 2016; Pérez-Montarelo et al., 2018; Grundstad et al., 2019). In USA100 *S. aureus* isolates (classified into CC5), the Agr system was confirmed to regulate production of hemolysins and proteases and to promote virulence in a murine model of skin infection (Grundstad et al., 2019). Much remains unknown regarding the pathogenesis of *S. aureus* CC5, in particular as it relates to IE.

In the present study, we used 8 methicillin-sensitive *S. aureus*, invasive isolates, classified into CC5 to investigate the association of IE with their expression of six global regulators of virulence (*RNAIII* [*agr*], *rot*, *sarS*, *sarA*, *sigB*, and *mgrA*). We found that vegetation formation, as tested in the rabbit model of left-sided, native valve IE, inversely correlates with *RNAIII* and *sarA* expression during growth in TH (beef heart infusion) broth. In fact, a strain deficient in *RNAIII* expression produced some of the largest vegetations, indicating that the *RNAIII*-regulated virulence genes encoded by the strain are not required for vegetation formation on native valves. However, this strain was deficient in causing systemic pathology and lethal sepsis. We also provided evidence that large aortic valve vegetations accompanied by severe systemic toxicity arise from high level expression of colonization factors with a slower transition towards expression of the exotoxins. Simultaneous high expression of *RNAIII, sarA*, *sigB*, and *mgrA* leads to severe systemic toxicity but is the one phenotype assessed in this study that fails to promote vegetation formation.

## Materials and Methods

### Bacterial Strains and Growth Conditions

Two sets of clinical *S. aureus* isolates were obtained from a French national prospective multicenter cohort, VIRSTA, that were categorized into IE or BA groups (Bouchiat et al., 2015; Le Moing et al., 2015). Patients were categorized as having definitive IE as defined by the modified Duke criteria (Li et al., 2000). Patients that presented with negative trans-thoracic or trans-esophageal echocardiograms and did not meet post-hospital criteria for IE at the 3-month follow-up visit were defined as BA (Bouchiat et al., 2015; Le Moing et al., 2015). Staphylococcal strains were used from low-passage-number stocks. All staphylococcal strains were grown in beef heart infusion broth (Bacto™ Todd Hewitt, Becton Dickinson) at 37 °C with aeration (225 rpm) unless otherwise noted. Strains used in this study are listed in Table 1. For endocarditis experiments, strains were grown overnight and diluted and washed in phosphate buffered saline (PBS - 2 mM NaH_2_PO_4_, 5.7 mM Na_2_HPO_4_, 0.1 M NaCl, pH 7.4).

**Table 1.** Staphylococcus aureus strains tested from CC5 lineage

| Strain | Alias | Enterotoxin gene cluster | Classical enterotoxins | New enterotoxins |
| --- | --- | --- | --- | --- |
| ST2012 0206 | BA0206 | *seo, sem, sei, selu, sen, seg* |  | *sep, selx* |
| ST2012 2011 | BA0211 | *seo, sem, sei, selu, sen, seg* |  | *selx* |
| ST2011 1372 | BA1372 | *seo, sem, sei, selu, sen, seg* |  | *selx* |
| ST2010 1791 | BA1791 | *seo, sem, sei, selu, sen, seg* | *sed* | *ser, selj, selx* |
| ST2011 0560 | IE0560 | *seo, sem, sei, selu, sen, seg* | *sed* | *ser, selj, selx* |
| ST2010 1420 | IE1420 | *seo, sem, sei, selu, sen, seg* | *tstH, sec, sed* | *sel, ser, selj, selx* |
| ST2010 1789 | IE1789 | *seo, sem, sei, selu, sen, seg* |  | *selx* |
| ST2010 2295 | IE2295 | *seo, sem, sei, selu, sen, seg* | *tstH, sed* | *sel, sep, ser, selj, selx* |

### Superantigen Gene Screen

Genomic DNA was extracted from single colonies grown overnight on TSAII agar plates with 5% sheep blood (Becton Dickinson) using colony lysis solution (1% Triton-X100, 2 mM EDTA, 20 mM Tris-HCl pH 8.0) and incubating the lysate at 94 °C for 15 min. Amplification was carried out using Phusion HF DNA polymerase (New England Biolabs; NEB) according to manufacturer’s instructions with superantigen-specific primers (Salgado-Pabón et al., 2014). PCR annealing temperatures were calculated using Tm Calculator v1.12.0 (NEB).

### Rabbit Model of Native Valve, Left-Sided IE

The rabbit model of IE was performed as previously described (Salgado-Pabón et al., 2013). Briefly, New Zealand White Rabbits, male and female, weighing 2-3 kg were obtained from Bakkom Rabbitry (Red Wing, MN) and anesthetized with ketamine (dose range: 10-50 mg/kg) and xylazine (dose range: 2.5-10 mg/kg). Mechanical damage to the aortic valve was done by introducing a hard plastic catheter via the left carotid artery, left to pulse against the valve for 2 h, removed, and the incision closed. Rabbits were inoculated via the marginal ear vein with 1.3 – 3.6×10^7^ cfu in PBS and monitored 4 times daily for failure to right themselves and failure to exhibit escape behavior. Simultaneous presence of these criteria is 100% predictive of fatal outcome and represents a humane endpoint. Infection was allowed to proceed for a total of 4 days unless a humane endpoint point was reached. For pain management, rabbits received buprenorphine (dose range: 0.01 – 0.05 mg/kg) twice daily throughout the duration of the experiment. At the conclusion of each experiment, venous blood was drawn and plated onto TSA II agar plates with 5% sheep blood for bacterial counts (Becton Dickinson). Rabbits were euthanized with Euthasol (Virbac) and necropsies performed to assess overall health. Spleens were weighed, kidney pathology was graded using a gross lesion pathology scale (Table S1), aortic valves were exposed to assess vegetation growth, and vegetations that formed were excised, weighed, and suspended in PBS for bacterial counts. Gross pathology grading was developed by a board-certified veterinary pathologist and done in a blinded manner (Gibson-Corley et al., 2013). All experiments were performed according to established guidelines and the protocol approved by the University of Iowa Institutional Animal Care and Use Committee (Protocol 6121907). All rabbit experimental data is a result of at least 2 independent experiments per infection group.

### Erythrocyte Lysis Assays for Hemolysin Production

Erythrocyte lysis assays were performed as previously described (King et al., 2016). Overnight cultures were diluted to an OD_600_ 1.0 in PBS and 5 µl spotted onto TSA II agar plates with either 5% rabbit or sheep blood (Becton Dickinson). Plates were incubated for 24 h at 37 °C with 5% CO_2_. Zones of hemolysis were measured and quantified using ImageJ. Data is represented by three independent experiments done with technical duplicates.

### Construction of qPCR Standard Curve Template Plasmid

qPCR primers for staphylococcal superantigens and virulence factor regulators were created using the PrimerQuest Tool from Integrated DNA Technologies (IDT) (Table S2). *S. aureus* strains MW2, MN8, N315, and IA209 were used as template sequences. A gBlock from IDT was ordered with all *S. aureus* SAg and regulator target amplicons listed in Table S2. The gBlock was inserted into a BamHI linearized pUC19 vector by Gibson Assembly creating pKK81 (Gibson et al., 2009). The resultant plasmid was verified by Sanger sequencing. All qPCR standard curves were made using SacI linearized pKK81 template and diluted with IDTE pH 8.0 buffer supplemented with 0.1 mg/mL tRNA (IDT) in DNA LoBind microfuge tubes (Eppendorf).

Plasmid concentration was determined using the NanoDrop™ 2000c (Thermo Fisher). Concentration was converted to template copy number by using the following equation: (*C*)(6.0221 × 10^23^ molecules/mole)/(N × 660 g/mole)(1 × 10^9^ ng/g) = copy number/µL (IDT). *C* is the concentration of template in ng/µL, N is the length of the dsDNA amplicon, and 660 g/mole is the average mass of 1 bp of dsDNA.

### RNA Extraction

Overnight cultures of *S. aureus* strains listed in Table 1 were inoculated in beef heart infusion broth at an OD_600_ of 0.1 and grown at 37 °C with aeration (225 rpm). Isolates were collected for RNA extraction at OD_600_ values of 0.25-0.30 (3×10^8^ CFU/mL), 0.80-0.85 (8×10^8^ CFU/mL), 1.80-1.90 (18×10^8^ CFU/mL), and 4.80-5.0 (50×10^8^ CFU/mL). At each OD, 1×10^9^ CFUs were pelleted by centrifugation at 16,000 × g for 30 s. Supernatants were poured off, and pellets flash frozen in liquid nitrogen and stored at −80 °C for RNA extraction the next day. Bacterial pellets were resuspended in 650 µl of TRIzol (Invitrogen) and transferred to Powerbead Tubes (glass 0.1 mm) (Qiagen). Samples were placed in a FastPrep FP120 cell disrupter and run twice at full speed for 30 s with a 3-min incubation on ice between runs. An additional 350 µl of TRIzol was added to the samples and centrifuged at 16,000 × g for 60 s before being transferred to Phasemaker Tubes (Invitrogen). RNA isolation proceeded according to manufacturer’s instructions with 2 additional ethanol washes to remove any residual phenol or guanidine isothiocyanate. Contaminating gDNA was removed using the Turbo DNA-Free™ Kit (Invitrogen) with 6U of TURBO™ DNase. RNA quantity and purity were assessed using the NanoDrop™ 2000c (Thermo Fisher) with median A260/280 values of 1.93 (95% CI: 1.91-1.94) and A260/230 of 1.55 (95% CI: 1.42-1.63). RNA integrity was assessed for distinct 23s and 16s rRNA bands by non-denaturing gel electrophoresis. RNA (250 ng/sample) was added to 2X gel loading buffer II (95% formamide, 18 mM EDTA, 0.025% SDS, 0.025% xylene cyanol, and 0.025% bromophenol blue) and denatured at 95 °C for 5 min followed by rapid cooling on ice for 1 min. Samples were run on a 1.2% TBE gel at 100 V (10 V/cm length between electrodes) for 1 h.

### cDNA Synthesis and qPCR

cDNA synthesis was carried out using 200 ng of RNA with the High-Capacity cDNA Reverse Transcription Kit according to manufacturer’s instructions (Applied Biosystems). Each cDNA sample was diluted 1:5 (~2 ng/µL) in IDTE pH 8.0 buffer (IDT). qPCR was conducted using the 2X PrimeTime Gene Expression Master Mix (IDT) with each reaction having a final concentration of 1X PrimeTime Gene Expression Master Mix, 0.5X EvaGreen (Biotium), 500 nM forward and reverse primers (Table S2), 50 nM ROX (Thermo Scientific^TM^), and 1 µL of the template or diluted cDNA experimental sample in a final reaction volume of 10 µL. Reactions were run under fast-qPCR conditions recommended by IDT: Activation −95 °C 3 min, 40 cycle amplification/elongation −95 °C 5 s followed by 58 °C 30 s, Melt curve analysis −95 °C 5 s, 58 °C 30 s, and 95 °C 5 s. gDNA contamination from each sample was assessed using a total of 20 ng of RNA with reaction conditions as listed above with all samples having C_T_ values >35. No template controls were run with each qPCR plate and had C_T_ values >38. All qPCR reactions were run and analyzed using the standard curve quantitation method with a 10-fold dilution curve on the QuantStudio 3 Real-Time PCR System and analysis software v1.5.1 (Applied Biosystems) (Table S3). PCR efficiencies used for calculations are listed in Table S3. Each timepoint was collected in biological triplicates with each sample done in technical duplicate qPCR reactions.

### Statistical analyses

The Log-rank (Mantel Cox) test was used for statistical significance of survival curves. Two-way analysis of variance (ANOVA) was used to determine significance in gene expression changes throughout bacterial growth. For comparison of gene expression across mean areas under the curve (AUC), hemolytic production, spleen size, blood cfu/mL, and gross pathology grading, the one-way or two-way ANOVA with either the Holm-Šídák’s or Fisher’s LSD multiple comparisons tests were used for significance. Statistical significance of vegetation size across means was determined by one-way ANOVA Kruskal-Wallis test with uncorrected Dunn’s multiple comparison test. All statistical analyses were done using GraphPad Prism v9.3.1 software. *α* = 0.05.

## Results

### Superantigen profile of invasive *S. aureus* CC5 isolates

Of major importance is understanding the underlying mechanism that defines *S. aureus* potential for IE development and sequelae. For this purpose, we focused on *S. aureus* strains classified as clonal complex 5 (CC5) given their frequency in causing invasive disease in humans with severe complications, such as septic shock (Shorr et al., 2006; Wyllie et al., 2006). The invasive CC5 isolates in this study were randomly selected from a French national prospective multicenter cohort collected from patients with definitive IE (IE0560, IE1420, IE1789, IE2295) or those that did not meet the criteria for definitive infective endocarditis (IE), and hence classified as bacteremia (BA) isolates (BA0206, BA0211, BA1372, BA1791) (Li et al., 2000; Bouchiat et al., 2015; Le Moing et al., 2015). Having *S. aureus* isolates from endocarditis and bacteremia patients allows us to assess how intrinsic bacterial characteristics of invasive isolates correlate with IE development and/or disease severity, as tested in a rabbit model of left-sided, native valve IE.

Superantigens are critically important for *S. aureus* IE development and strains deficient in these virulence factors are significantly attenuated in experimental IE (Salgado-Pabón et al., 2013; Stach et al., 2016; Kinney et al., 2022). Given the contribution of superantigens to IE but knowing that they are variably encoded (Nienaber et al., 2011), we performed a genomic screen for the presence of 22 superantigens. *selX* and the enterotoxin gene cluster (*egc*) superantigens *seg, sei, sem, sen, seo,* and *selu* were detected in all 8 isolates (Table 1). *entC* (SEC gene) and *tstH* (TSST1 gene) were detected in only 2 of the isolates, both of IE origin. *entD* (SED gene), *sej*, *ser*, *sep*, and *sel* were variably present across isolates. *sea, entB* (SEB gene), *entE* (SEE gene), *seh, sek, seq, ses,* and *set* were not detected in any of the *S. aureus* isolates. Therefore, we confirmed that at minimum all the isolates encoded the *egc* superantigens, shown to be sufficient for IE development (Stach et al., 2016). Hence, all 8 strains have the potential to cause IE.

### Invasive *S. aureus* CC5 isolates vary in causation of IE and lethal sepsis

To establish the clinical manifestations of invasive *S. aureus* CC5 isolates in experimental IE, New Zealand White rabbits were injected intravenously with 1.3 – 3.6×10^7^, after 2 h mechanical damage to the aortic valve. Rabbits were monitored for up to 4 days. During that period, 5 isolates (BA0206, BA1791, IE0560, IE1420, and IE2295) formed similarly sized vegetations with median ranges of 20–32 mg (Fig. 1A). The largest vegetations were observed in BA0211 (median 82 mg) and IE1789 (median 94.5 mg) (Fig. 1A). The notable exception was BA1372, which was deficient in promoting IE with a vegetation median of 9.5 mg (Fig. 1A). As with vegetation formation, significant differences in lethality were directly related to individual strains. Most isolates (7/8) exhibited high lethality with less than 50% of rabbits surviving the infection (Fig. 1B-E). BA0206 and BA0211 were the most lethal with 100% of rabbits succumbing to the disease (Fig. 1B). In stark contrast, infection with IE1789 resulted in 83% survival during the experimental period despite being one of the strains producing the largest vegetations (Fig. 1A, E). Also of note is BA1372, which exhibited 50% lethality in spite of its deficiency in vegetation formation (Fig. 1A, C).

**Figure 1.**
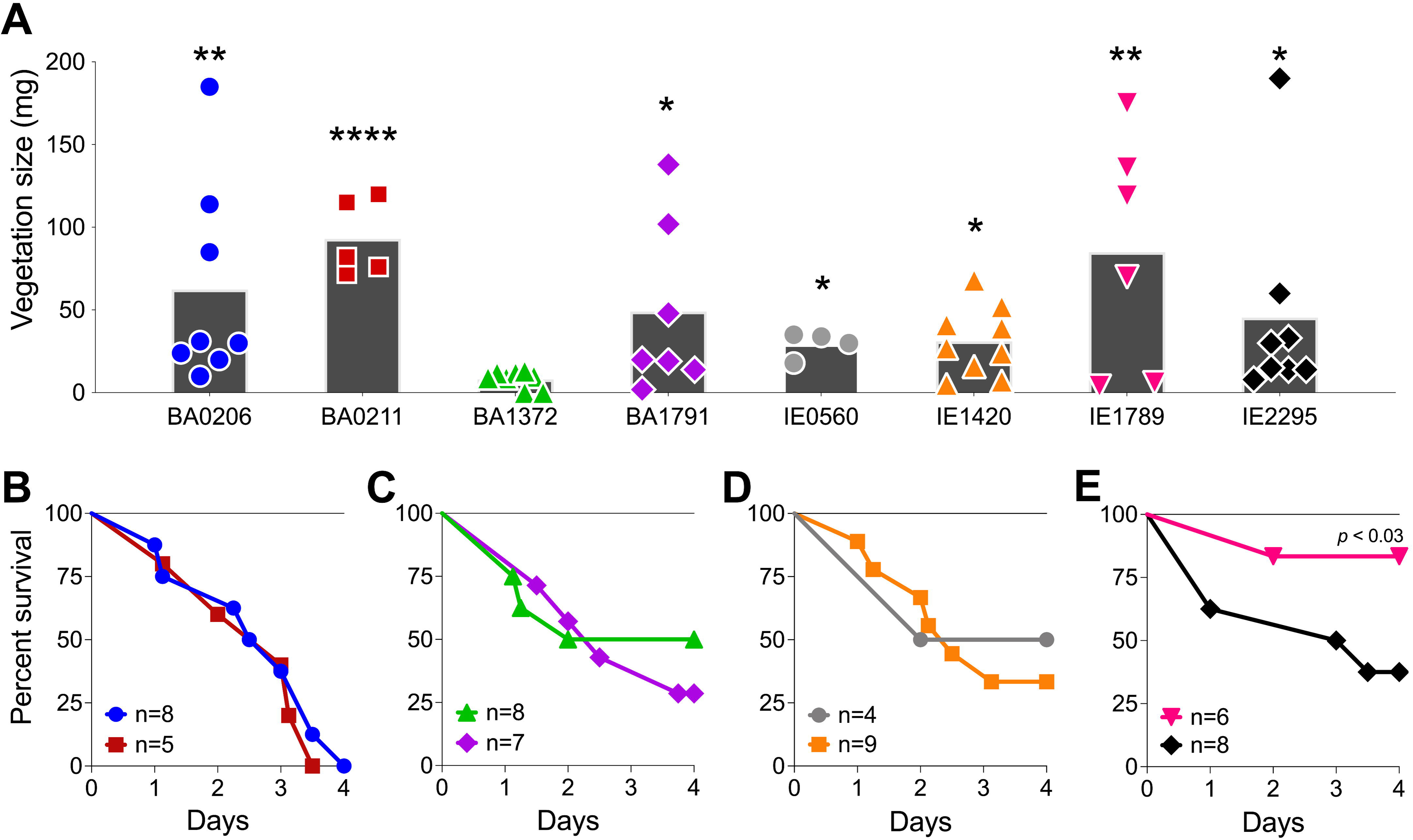
Differential development of infective endocarditis and lethality in *S. aureus* CC5 strains. Rabbit model of native valve IE and sepsis. Rabbits were injected intravenously with 1.3–3.6 × 10^7^ CFUs of *S. aureus* CC5 isolates after mechanical damage of aortic valve and monitored over 4 days. **(A)** Total weight of vegetations dissected from aortic valves. \**p* < 0.05, \*\**p* < 0.005, \*\*\*\**p* < 0.0001, one-way ANOVA, non-parametric Kruskal-Wallis with uncorrected Dunn’s multiple comparison test to BA1372 (IE-deficient). Bars represent mean value. **(B-E)** Percent survival. **(B)** BA0206 and BA0211. **(C)** BA1372 and BAA1791. **(D)** IE0560 and IE1420. **(E)** IE1789 and IE2295. *p <* 0.03, Log-rank (Mantel-Cox) of IE1789 compared to each strain.

*S. aureus* IE is commonly characterized by hematogenous spread with establishment of metastatic infections and systemic pathology (Tong et al., 2015). These complications also occur in our rabbit model and present as acute kidney injury, ischemic liver lesions, and lung injury. In this study, rabbits (n=55) were grossly assessed and scored for the presence of kidney lesion pathology on a scale from 0-3. Lesions in the *S. aureus* IE rabbit model present as hemorrhagic, necrotic, or ischemic (Fig. 2A). In the most severe pathology (grade 3), lesions are locally extensive, coalescing to diffuse, and extend across a large surface of the kidney (Table S1). Infection with *S. aureus* CC5 isolates led to severe pathology in >50% of rabbits (Fig. 2B). The severity of acute kidney injury was consistent with lethality exhibited by individual strains. BA0206 and BA0211 (strains with the highest lethality) induced grade 3 pathology in 100% of rabbits, while IE1789 (strain with the highest survival) induced no lesions or grade 1 pathology in >75% of rabbits (Fig. 2B). *S. aureus* isolates were recovered from the bloodstream at endpoints in the range of 1×10^3^ – 1.3×10^5^ cfu/mL (Fig. 2C), with the sole exception being IE1789 where fewer than 100 bacteria were detected in the bloodstream in 5/6 rabbits (Fig. 2C). The data indicates that most (5/8) strains have common characteristics that drive IE development and systemic complications to similar extent. However, it is also evident that individual strain characteristics determine the severity of the disease.

**Figure 2.**
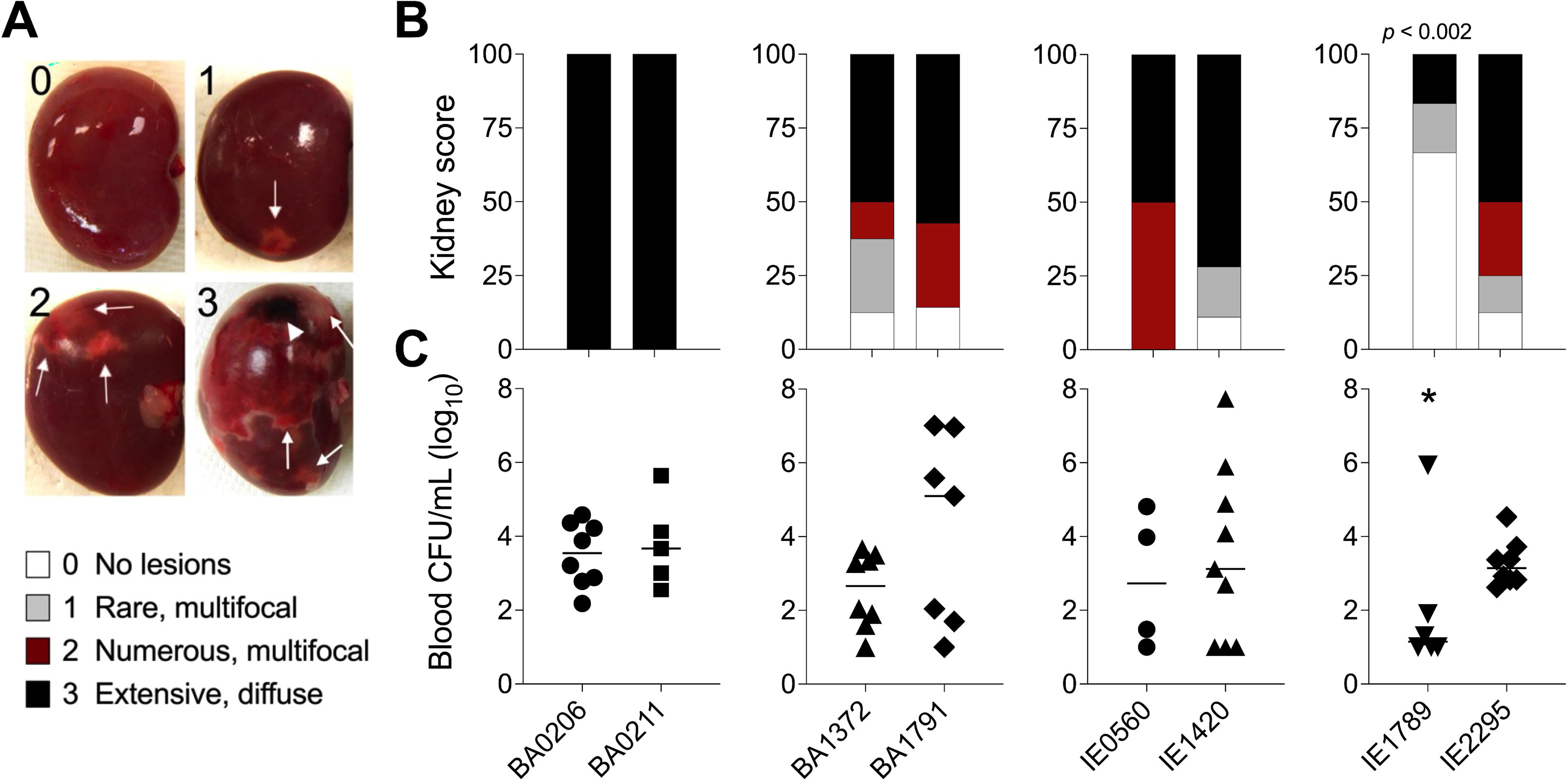
Acute kidney injury and bacteremia resulting from *S. aureus* CC5 IE. Rabbit model of native valve IE and sepsis. Rabbits were injected intravenously with 1.3–3.6 × 10^7^ cfu of *S. aureus* CC5 isolates after mechanical damage of aortic valve and monitored over 4 days. (**A**) Kidney gross pathology grading scale (grades 0-3). 0 = no lesions, 1 = rare, small (<4mm) multifocal lesions, 2 = numerous large (>5mm) multifocal lesions, 3 = extensive to coalescing to diffuse lesions. Arrows indicate ischemic and/or hemorrhagic lesions, arrowhead indicates a necrotic lesion. (**B**) Scoring of kidney lesions *post-mortem*. One-way ANOVA with Fisher’s LSD multiple comparisons test across strains. *p* < 0.002, IE1789 compared to the rest of the strains. (**C**) Bacterial counts per milliliter of blood recovered from rabbits *post-mortem*. Lines represent median value. *, *p* < 0.02, one-way ANOVA Kruskal-Wallis test with uncorrected Dunn’s multiple comparison test across strains.

Extensive phenotypic variation among clinical isolates is a growing subject in medical microbiology (Jelsbak et al., 2010). In *S. aureus*, major strain-dependent differences in gene expression are in part caused by differential expression of global regulators and two-component systems (Jelsbak et al., 2010; Zhao et al., 2019). *RNAIII*, *rot*, *sarA*, *sarS*, *sigB*, and *mgrA* have previously been shown to influence expression of *S. aureus* virulence factors (Schmidt et al., 2004; Ingavale et al., 2005; Tseng and Stewart, 2005; Andrey et al., 2010, 2015; Kusch et al., 2011; Tuffs et al., 2019). Hence, we addressed their gene expression in our strain collection next.

### Differential expression of *RNAIII* and *rot* in CC5 strains

RNAIII is a regulatory mRNA and effector molecule of the Agr quorum-sensing system that differentially controls expression of *S. aureus* surface proteins [e.g. microbial surface components recognizing adhesive matrix molecules (MSCRAMMs) and protein A] and the secreted toxins and enzymes (e.g. hemolysins, superantigens, proteases, nucleases, lipases) (Jenul and Horswill, 2019). *RNAIII* expression itself is regulated by the Agr system. When the Agr system is active, RNAIII is expressed which in turn represses surface proteins and induces exoproteins. The opposite is also true. RNAIII control of exoprotein production is achieved indirectly via translational suppression of the global regulator Rot (Mcnamara et al., 2000; Geisinger et al., 2006). Rot (for *repressor of toxins*) inhibits transcription of toxins and extracellular proteases during growth in exponential phase, when the Agr system is inactive and RNAIII is uninduced, while at the same time induces production of ClfA, coagulase, protein A, and the transcriptional regulator SarS (Saïd-Salim et al., 2003) (Fig. S1).

Traditionally, comparative C_T_ quantitation is performed for studies of gene expression. The biggest challenge with this method is comparing and interpreting relative expression data between studies that use non-isogenic strains or strains with phenotypic plasticity, as deviations in expression of the housekeeping gene(s) of choice changes the results (Pfaffl, 2004; Huggett et al., 2005; Valihrach and Demnerova, 2012). This is the case for the *S. aureus* CC5 collection. We quantified absolute copies of the *gyrB* transcript, a common *S. aureus* housekeeping gene (Valihrach and Demnerova, 2012; Sihto et al., 2014; Crosby et al., 2016; Stach et al., 2016), by RT-qPCR using the standard curve method (Fig. S2) (Bustin, 2000; Pfaffl, 2004). For this purpose, *S. aureus* isolates were grown to specific cell densities for a period of 6 h in TH broth. Growth in exponential phase (2.8 – 18×10^8^ cfu/mL) resulted in stable *gyrB* expression (ΔC_T_ < 0.5) in 7 out of 8 isolates. IE2295 was the only strain with a 2-fold decrease in *gyrB* expression (ΔC_T_ > 0.5) during exponential growth (Fig. S2B). Yet, *gyrB* expression was not stable during growth in stationary phase (18 – 50×10^8^ cfu/mL) with half the strains exhibiting 2-fold decreases that could potentially affect experimental outcomes (Fig. S2) (Bustin, 2000; Pfaffl, 2004; Huggett et al., 2005; Valihrach and Demnerova, 2012). Therefore, we favored absolute RT-qPCR analysis as a method better suited to address expression levels of multiple genes within strains and across different strains.

In our *S. aureus* CC5 collection of invasive isolates, *RNAIII* expression increased 10 – 3000-fold from early to post-exponential growth in 7 out 8 of strains (Fig. 3A, B). IE1789 and IE2295 started with less than 1000 copies/ng of RNAIII, but while in IE2295 *RNAIII* expression was rapidly induced to more than a million copies/ng, *RNAIII* expression in IE1789 remained low and uninduced (Fig. 3B). BA0206 and IE0560 started with the highest RNAIII levels but only expression in BA0206 continued to increase through growth in stationary phase (Fig. 3A, B) resulting in the strain having the highest RNAIII levels in the collection (Fig. 3C). Three other strains also exhibited significantly higher levels of RNAIII, BA1372, BA1791, and IE2295 (Fig. 3C). BA0211 and IE1420 were unique in that *RNAIII* expression significantly increased during post-exponential growth reaching close to top-level expression at 50×10^8^ cfu/mL (Fig. 3A, B).

**Figure 3.**
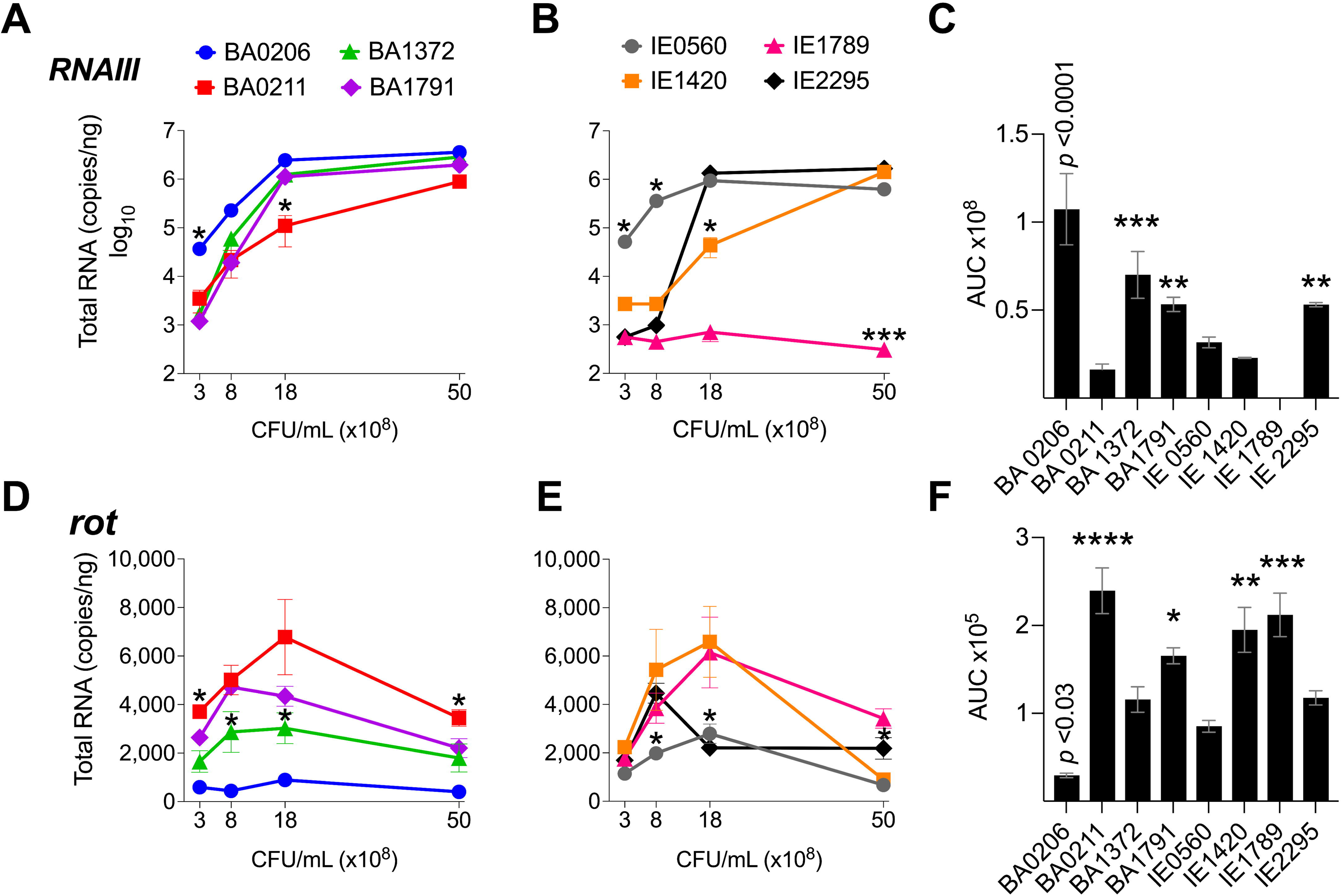
*RNAIII* and *rot* expression in *S. aureus* CC5 isolates. Quantitation of *S. aureus* CC5 gene expression during growth in TH broth by RT-qPCR standard curve quantitation method. **(A-B)** *RNAIII* expression and **(D-E)** *rot* expression at indicated cell densities. Error bars (standard deviation) not shown are smaller than symbol. Asterisks indicate data points significantly different than the rest at that specific cell density. **(C, F)** Area under the curve (mean ± SEM). Data is the result of three biological replicates. *, *p* < 0.05, **, *p* < 0.005, ***, *p* < 0.0005, ****, *p* < 0.0001, one-way ANOVA with Holm-Šídák’s multiple comparisons test across strains.

*rot* expression was inversely proportional to *RNAIII* expression, peaking at 8 – 18×10^8^ cfu/mL (Fig. 3D-F). In most strains, *rot* expression decreased during post-exponential growth concomitant to *RNAIII* expression (Fig. 3D, E). IE1789 (the lowest *RNAIII* expressor) expressed *rot* at high levels while BA0206 (the highest *RNAIII* expressor) exhibited low and uninduced *rot* expression (Fig. 3D, F). Given that expression of most *S. aureus* hemolysins is controlled by RNAIII and Rot, we tested the hemolytic activity of the CC5 isolates against both rabbit and sheep erythrocytes. As expected, IE1789 was non-hemolytic against rabbit erythrocytes (Fig. 4A) and showed very low hemolytic activity against sheep erythrocytes (Fig. 4B). The rest of the strains exhibited hemolytic activity consistent with expression of hemolysins such as *α*-toxin, *β*-toxin, and phenol-soluble modulins (PSMs) (Fig. 4). From these results, it becomes evident that *RNAIII* expression is not a critical requirement for *S. aureus* vegetation formation, as exemplified by strains IE1789 (proficient) and BA1372 (deficient).

**Figure 4.**
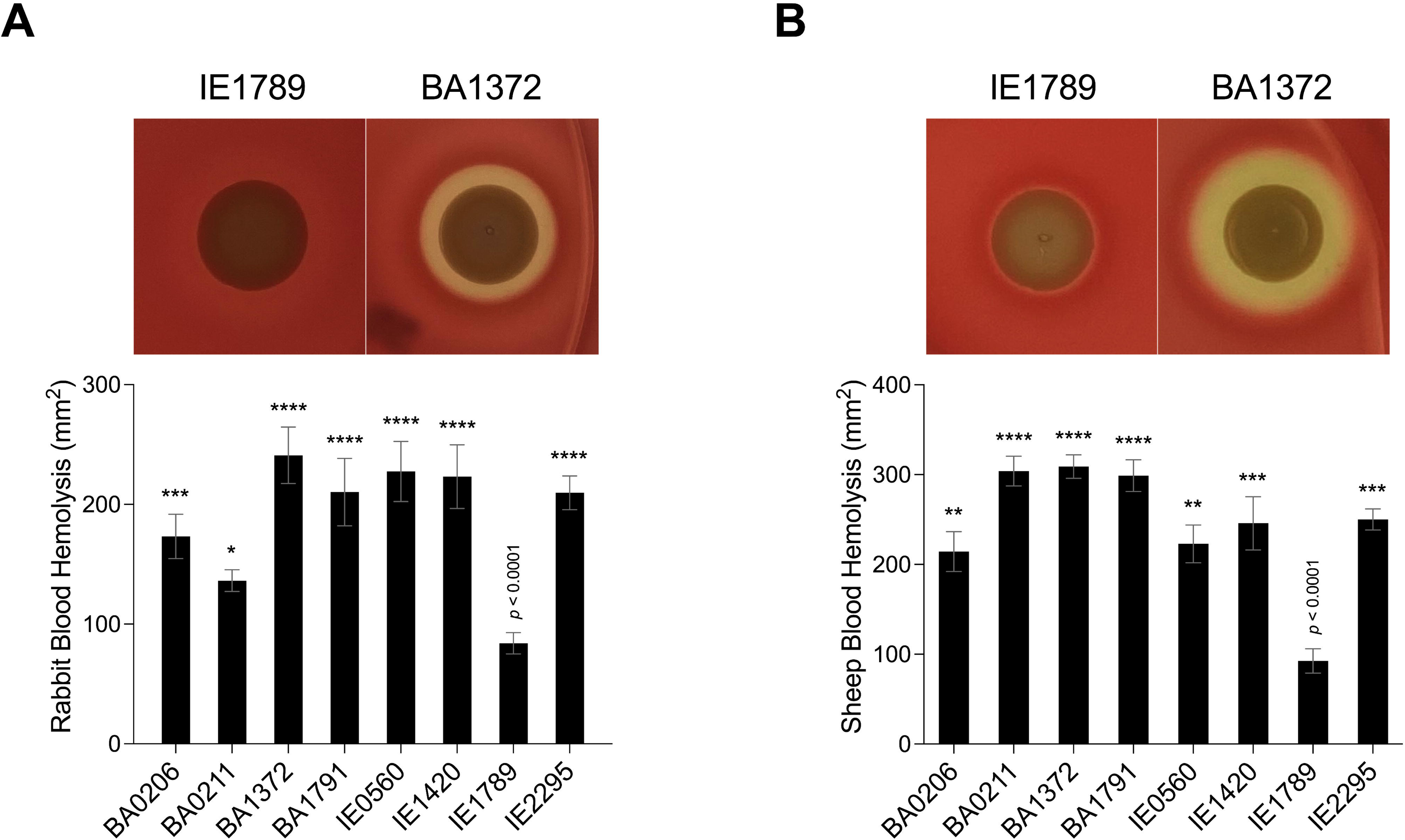
Hemolytic activity in *S. aureus* CC5 isolates. Overnight cultures of *S. aureus* CC5 isolates were washed and spotted onto **(A)** 5% rabbit blood and (**B**) 5% sheep blood TSA II agar plates. Zones of hemolysis were measured after overnight growth. (Top) Representative images of hemolytic activity in strains IE1789 and BA1372. (Bottom) Relative levels of hemolysin production as measured in an erythrocyte lysis assay. Data are represented as mean ± SEM. *, *p* < 0.05, **, *p* < 0.005, ***, *p* < 0.0005, ****, *p* < 0.0001, one-way ANOVA with Holm-Šídák’s multiple comparisons test to the non-hemolytic strain IE1789.

### Differential expression of *sarA* and *sarS* in CC5 strains

SarA is known to increase expression of the *agr* system (Heinrichs et al., 1996; Rechtin et al., 1999; Jenul and Horswill, 2019) and to repress expression of the transcriptional regulators *sarS* and *rot* (Cheung et al., 2001; Hsieh et al., 2008) and genes encoding for protein A and proteases (Morrison et al., 2012). SarS acts opposite to SarA by inducing *spa* (Protein A) gene expression and repressing expression of several toxin genes such as *hla* (*α*-toxin gene) (Fig. S1) (Tegmark et al., 2000). *sarA* expression patterns were more diverse among the *S. aureus* CC5 isolates. During exponential growth, *sarA* was significantly induced in strains BA206 and IE0560, while repressed in IE1789 (Fig. 5A, B). During post-exponential growth, *sarA* was significantly induced in BA1372, while repressed in BA1791, IE0560, and IE1420 (Fig. 5A, B). BA0211 was the only strain that did not exhibit significant *sarA* induction throughout growth (Fig. 5A, B). Overall, BA1372, IE0560, and IE2295 exhibited the highest sarA levels (Fig. 5C). Consistent with the literature, *sarA* expression was inversely proportional to *sarS* (Fig. 5D-F). The sole exception being BA206, which exhibited low and uninduced expression of *sarS*, not concomitant with *sarA* expression (Fig. 5D) but consistent with low and uninduced *rot* (Fig. 2D). Significantly higher levels of sarS were seen in BA0211, BA1791, IE1420, and IE1789 (Fig. 5F). These strains also exhibited higher levels of rot, a positive regulator of sarS expression (Fig. 2F) (Saïd-Salim et al., 2003). No requirement was evident for *sarA* induction in vegetation formation, as exemplified by strains BA0211 and IE1789.

**Figure 5.**
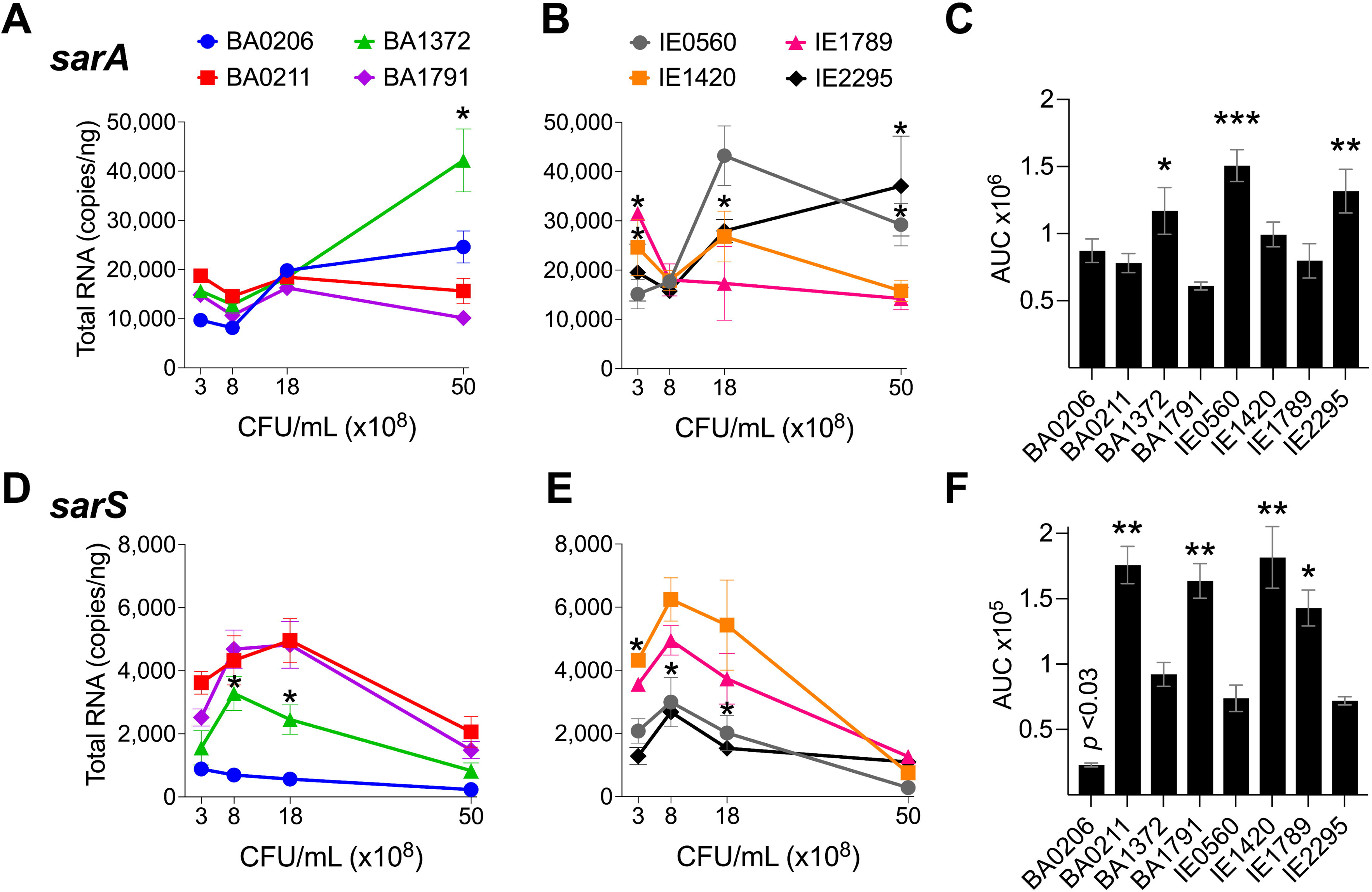
*sarA* and *sarS* expression in *S. aureus* CC5 isolates. Figure 3. Quantitation of *S. aureus* CC5 gene expression during growth in TH broth by RT-qPCR standard curve quantitation method. **(A-B)** *sarA* expression and **(D-E)** *sarS* expression at indicated cell densities. Error bars (standard deviation) not shown are smaller than symbol. Asterisks indicate data points significantly different than the rest at a specific cell density. **(C, F)** Area under the curve (mean ± SEM). Data is the result of three biological replicates. *, *p* < 0.05, **, *p* < 0.005, ***, *p* < 0.0005, one-way ANOVA with Holm-Šídák’s multiple comparisons test across strains.

### Differential expression of *sigB* and *mgrA* in CC5 strains

The alternative sigma factor B (SigB) controls hundreds of genes, many of which are involved in stress responses as well as virulence (Guldimann et al., 2016). SigB directly inhibits expression of the *agr* operon (Bischoff et al., 2001), but induces expression of *sarA*, adhesin genes (e.g ClfA and FnBPs genes), and the *egc* superantigen operon (Bischoff et al., 2004; Entenza et al., 2005; Kusch et al., 2011). SigB is also reported to represses expression of *rot* specifically during growth in stationary phase (Fig. S1) (Hsieh et al., 2008). In our collection of invasive *S. aureus* strains, *sigB* expression peaked early at 8×10^8^ cfu/mL but was followed by a rapid drop in expression with further growth (Fig. 6A, B). Of the isolates with the highest *sigB* expression (Fig. 6C), BA0206, BA0211, and BA1372 achieved it by maintaining expression in post-exponential growth (Fig. 6A), while IE1420 and IE1789 achieved it by inducing it at high levels in exponential phase (Fig. 6B). It was of interest to establish whether the temporal expression of *sigB* in high-expressing isolates could result in differential gene expression of *sigB-*regulated genes important for IE development. For that purpose, we tested BA1372 (IE deficient) and IE1789 (IE proficient) for expression of the *egc* operon, which promotes vegetation formation and is induced by SigB (Kusch et al., 2011; Stach et al., 2016). In IE1789, *egc* expression was highest during exponential growth (Fig. 7A) whereas in BA1372 *egc* expression remained low throughout (Fig. 7B). Consistent with SigB induction of *egc* superantigens specifically, expression of *selX* (reported to be under *saeRS* control) was almost identical in both strains (Fig. 7A, B) (Langley et al., 2017). Hence, temporal expression of master regulators contributes to the phenotypic heterogeneity of *S. aureus* strains and possibly variable disease outcomes.

**Figure 6.**
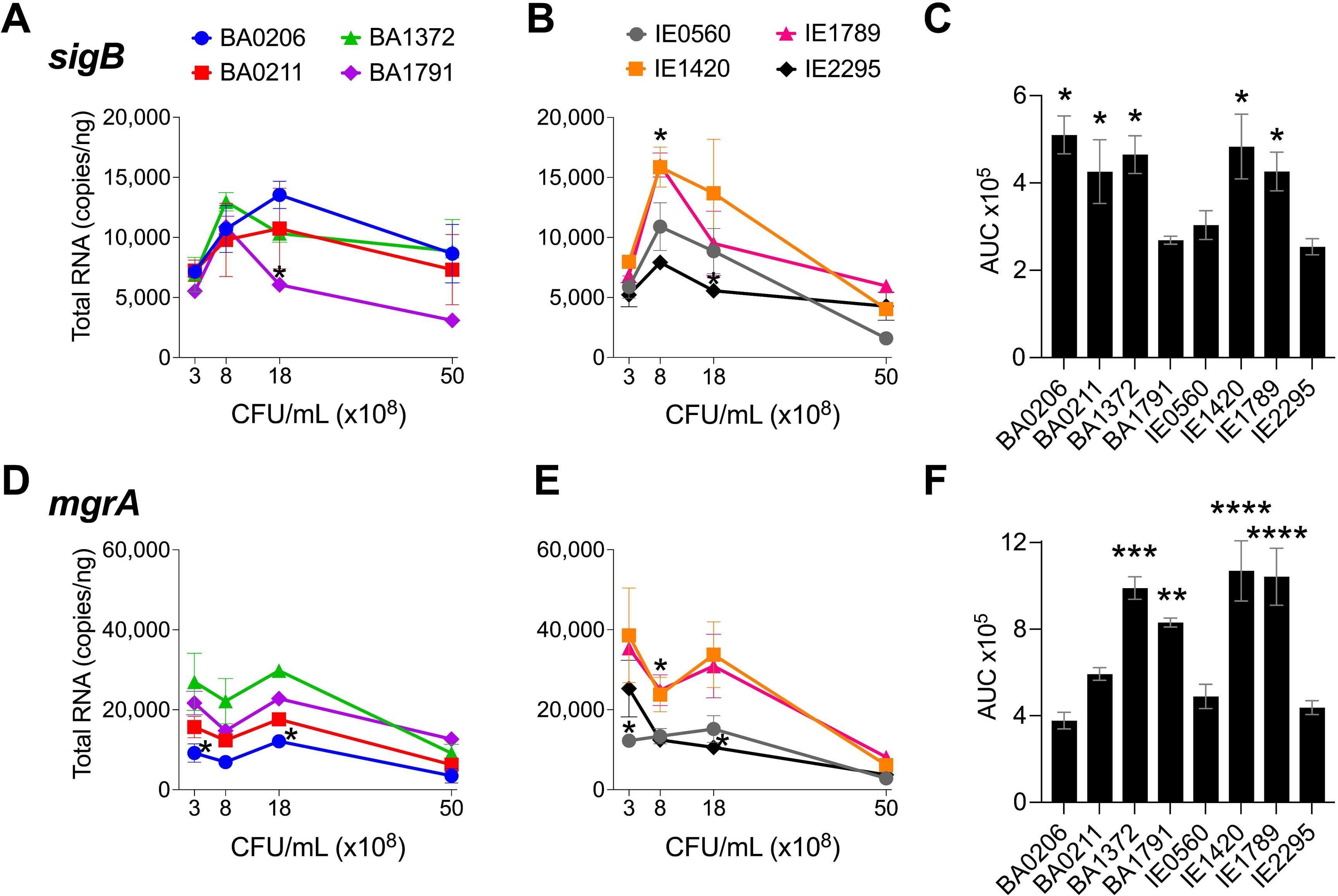
*sigB* and *mgrA* expression in *S. aureus* CC5 isolates. Quantitation of *S. aureus* CC5 gene expression during growth in TH broth by RT-qPCR standard curve quantitation method. **(A-B)** *sigB* expression and **(D-E)** *mgrA* expression at indicated cell densities. Error bars (standard deviation) not shown are smaller than symbol. Asterisks indicate data points significantly different than the rest at a specific cell density. **(C, F)** Area under the curve (mean ± SEM). Data is the result of three biological replicates. *, *p* < 0.05, **, *p* < 0.005, ***, *p* < 0.0005, one-way ANOVA with Holm-Šídák’s multiple comparisons test across strains.

**Figure 7.**
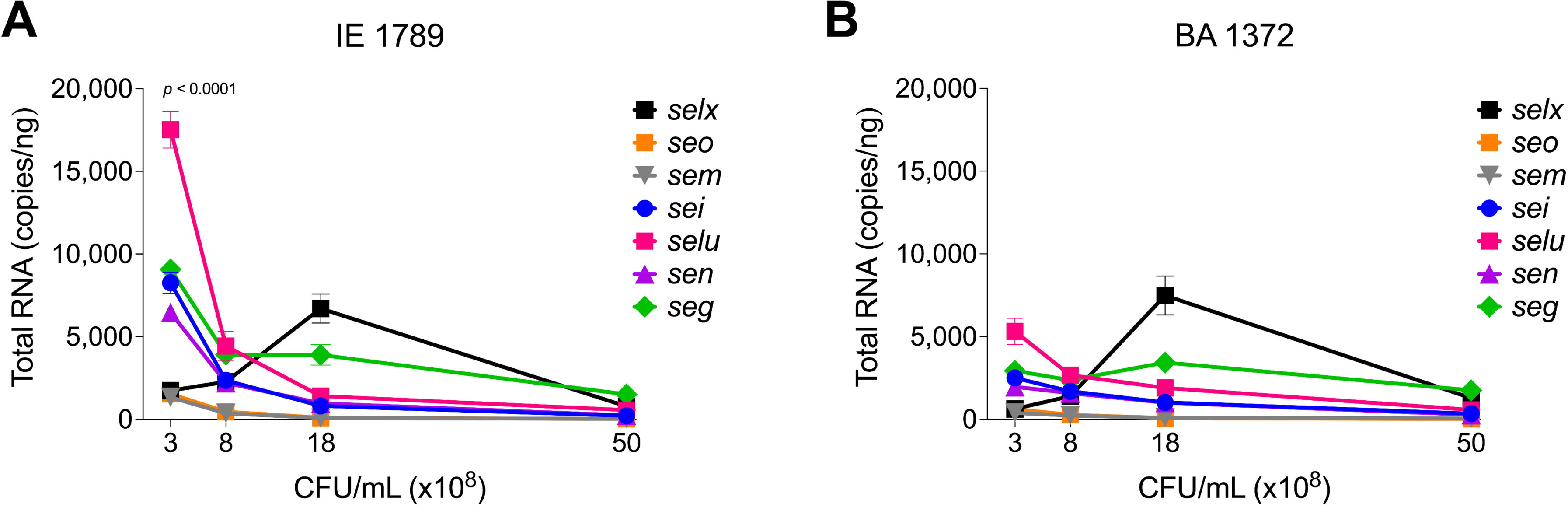
Differential expression of *egc* SAgs and *selx*. Quantitation of egc mRNA in *S. aureus* CC5 during growth in TH broth by RT-qPCR standard curve quantitation method. *egc* and *selx* expression in **(A)** IE1789 and **(B)** BA1372 at indicated cell densities. Error bars (standard deviation) not shown are smaller than symbol. Data is the result of three biological replicates. Two-way ANOVA with Holm-Šídák’s multiple comparisons test across superantigen pairs in IE1789 and BA1372 during exponential growth.

MgrA regulatory effects mirror those of the *agr* system, although the specific targets can be distinct (Luong et al., 2006). MgrA upregulates production of secreted proteins (e.g. leukotoxins, Spl proteases, enzymes) and downregulates production of surface-associated proteins (e.g. large surface protein Ebh, SraP, SasG; Fig. S1) (Luong et al., 2006; Crosby et al., 2016). Initial mgrA levels in our *S. aureus* CC5 collection ranged from 10,000 – 40,000 copies/ng and remained stable and at its highest throughout exponential growth (Fig. 6D, E). Once in post-exponential growth, *mgrA* expression significantly decreased in all strains. The one exception was IE2295 which exhibited a significant decrease in *mgrA* expression during exponential growth (Fig. 6E). Half the strains expressed *mgrA* at significantly higher levels with a more distinct separation of high versus low expressors within the endocarditis isolates (Fig. 6F).

### Inverse correlation between *RNAIII* and *sarA* expression and vegetation formation in *S. aureus* CC5 strains

A remaining question is, what drives differential vegetation formation in these isolates? To address this question, we performed a Pearson correlation analysis of gene expression among the six global regulators examined in this study and median vegetation size produced by the eight *S. aureus* CC5 isolates. Initial clustering of the CC5 strains based on their clinical manifestation (bacteremia or endocarditis) was insightful as it provided evidence of divergent expression patterns of some of the bacteremia strains tested herein (Fig. 8A). Consistent with the literature, *RNAIII* expression inversely correlated with *rot*, and *rot* expression directly correlated with *sarS* in all isolates. *sarA* expression inversely correlated with *sarS* in 7/8 isolates (exception BA0206), but it only correlated with *RNAIII* expression in 5/8 strains (exceptions BA0206, BA1791, and IE0560). The most striking difference across isolates was the correlations between *sigB* expression and the rest of the regulators examined in this study (Fig. 8A). In this context, for every correlation observed within the *S. aureus* CC5 collection, 2-3 of the bacteremia isolates were the exception. *sigB* expression inversely correlated with *RNAIII* in 6/8 strains (exceptions BA206 and BA1372) and with *sarA* in 6/8 strains (exceptions BA1372 and BA1791), while it directly correlated with *rot* in 5/8 strains (exceptions BA0206, BA1372, and BA1791) and with *mgrA* in 5/8 strains (exceptions BA0206, BA0211, and BA1791) (Fig. 8A). The inverse correlation between *sigB* and *sarA* expression in most of the CC5 isolates in our collection was surprising given that SigB is reported to upregulate SarA (Manna et al., 1998).

**Figure 8.**
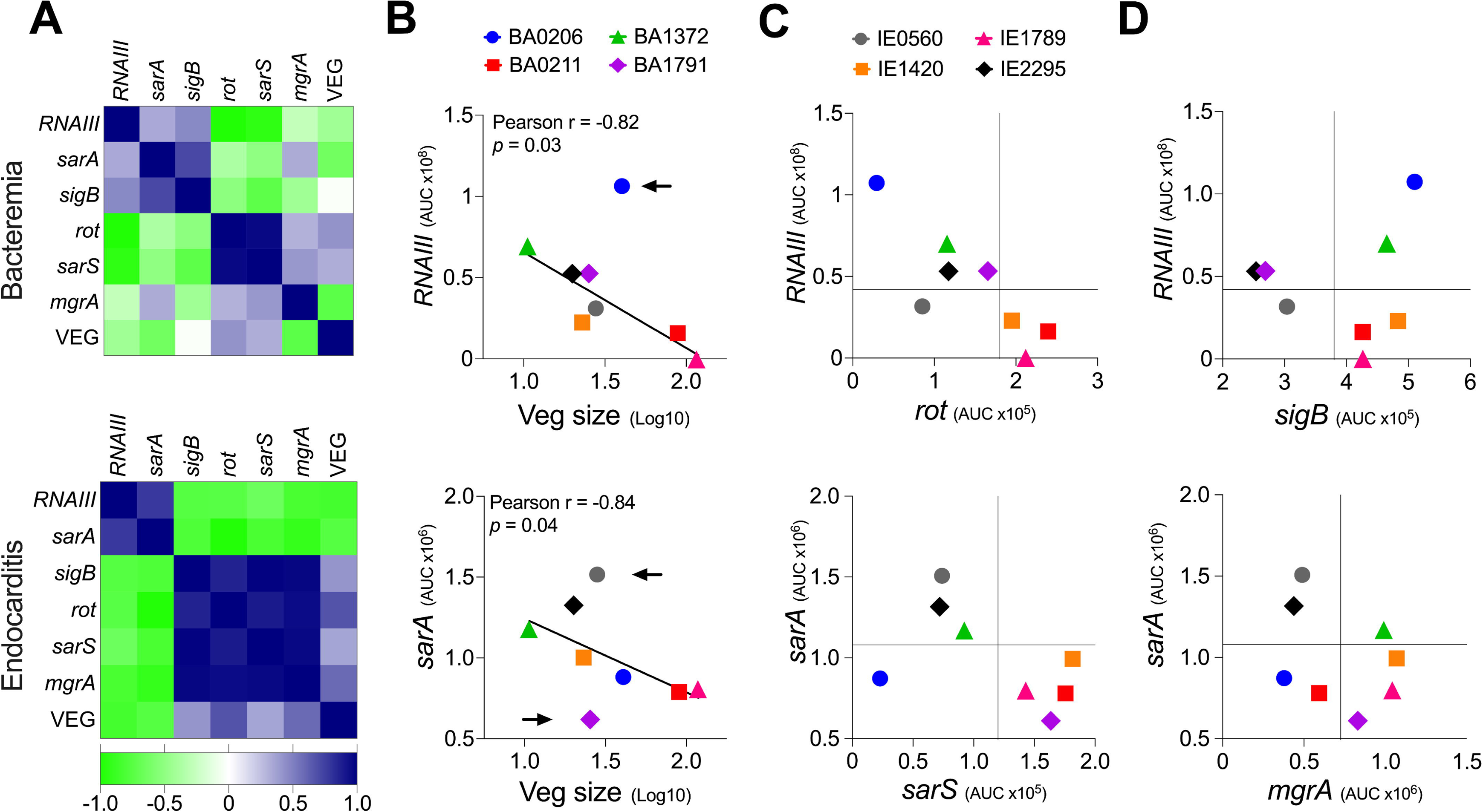
Vegetation formation inversely correlates with *RNAIII* and *sarA* expression in *S. aureus* CC5 isolates. **(A)** Heat map of correlation matrix of gene expression across regulators (expressed as AUC) and vegetation size. Blue, direct correlation of expression. Green, inverse correlation of expression. **(B)** Correlation between vegetation size and *RNAIII* expression (top panel) or *sarA* expression (bottom panel). Pearson correlation coefficients (Pearson r), correlation *p* value, and best-fit line shown. Arrows indicate points that fall outside the trend shown. **(C-D)** Distribution of expression patterns across strains (expressed as AUC). Perpendicular lines mark where statistical significance in expression is achieved for the regulators shown in the graph.

Of great interest, vegetation size inversely correlated with both *RNAIII* expression and *sarA* expression, with a stronger correlation observed with *RNAIII* (Fig 8A, B). It was intriguing that BA0206 produced large vegetations despite having the highest *RNAIII* expression, in particular when BA1372 was deficient in vegetation formation. To address this, we analyzed the expression patterns of the six regulators studied herein across all eight strains. In BA0206, *RNAIII* expression was high yet *sarA* expression was low (Fig 8B). High *RNAIII* expression was accompanied by concomitantly low *rot* and *sarS* expression (Fig. 8C) and unconventionally by high *sigB* expression (Fig. 8D). BA1372 expressed *RNAIII* and *sarA* at high levels (Fig 8B) in addition to expressing *sigB* and *mgrA* at high levels (Fig. 8D). Altogether, the combination of high expression of *RNAIII*, *sarA*, *sigB*, and *mgrA* was unique to BA1372 in our collection, while the combination of high *RNAIII* and *sigB* with low *sarA* and *mgrA* was unique to BA0206. Overall, these results highlight the heterogeneity among the CC5 lineage but reveals expression patterns specific to IE deficient and proficient strains.

## Discussion

*S. aureus* CC5 isolates are prevalent colonizers and agents of infection in the U.S. and have become a predominant IE clonal group (Enright et al., 2002; Tenover et al., 2008; Limbago et al., 2009; Roberts, 2013). However, it has been long recognized that *S. aureus* exists as a heterogenous population showing extensive phenotypic variation with regulation of single genes that can vary considerably even within clonal groups (Jelsbak et al., 2010). Regulation of virulence factors results from a complex network of host and environmental cues that elicit a coordinated response. Changes in the levels of major components of these regulatory networks affects how *S. aureus* responds to a given cue. In the present study, we investigated the temporal expression of prominent global regulators of virulence (*RNAIII* [*agr*], *rot*, *sarS*, *sarA*, *sigB*, and *mgrA*) in eight invasive *S. aureus* CC5 isolates and established intrinsic expression patterns associated with IE outcomes. We provide evidence that vegetation formation, as tested in the rabbit model of left-sided native valve IE, inversely correlates with *RNAIII* and *sarA* expression when grown in beef heart infusion broth.

Of interest, even with a small collection of isolates from the same clonal group, 4 distinct clinical outcomes were observed: (i) strains with similar vegetation size, kidney injury, and lethality (BA1791, IE0560, IE1420, and IE2295), (ii) a strain that produces average-size vegetations but causes more severe kidney injury with increased lethality (BA0206), (iii) strains that produce significantly larger vegetations but differ in potential to cause kidney injury and lethal sepsis (IE1789 and BA0211), and (iv) a strain that is deficient in vegetation formation but proficient in causing kidney injury and lethal sepsis (BA1372).

Based on the correlation analysis, one would have expected BA0206 to be less efficient at promoting IE, as it exhibits the highest *RNAIII* expression concomitant with basal levels of rot and sarS transcript. Yet, BA0206 is one of five strains that are similar in their ability to promote vegetation formation. Furthermore, infection with BA0206 results in dissemination, systemic toxicity, and lethality indistinguishable from that of BA0211 (strain that produces exceptionally large vegetations and lethal sepsis). While *RNAIII* expression is high, this is not reflected in its hemolytic activity against both rabbit and sheep erythrocytes, which together are sensitive to the action of the pore-forming toxins, phenol-soluble modulins, and the sphingomyelinase *β*-toxin. The lower *sarA* expression levels in this strain may help counter-balance high expression of *RNAIII*. Low SarA increases expression of proteases that degrade extracellular virulence factors, which can help explain the lower hemolytic activity (Zielinska et al., 2012). High *sigB* expression (surprising for a strain with high *RNAIII* expression) can induce production of colonization factors required for vegetation formation.

IE1789 and BA0211 are highly proficient in forming large aortic valve vegetations yet induce drastically divergent systemic outcomes. IE1789 has no inducible expression of *RNAIII*, resulting in no or minimal hemolysis. High expression of *rot, sarS,* and *sigB* indicates that IE1789 is locked in a colonization state as these regulators are known to induce production of MSCRAMMs (e.g. clumping factors, fibronectin-binding proteins, protein A), enzymes (e.g. coagulase), and the *egc* superantigens. Many of these virulence factors have been shown to contribute to development of *S. aureus* IE in experimental models (Vernachio et al., 2003; Que et al., 2005; Malachowa et al., 2011; McAdow et al., 2011; Stach et al., 2016). These results provide further evidence that the Agr-regulated exoproteins (at minimum the pore-forming toxins) are not critical for establishing *S. aureus* IE and provide insights into the minimal requirements for vegetation formation on native valves. Similar observations have been reported with *S. aureus agr* mutants tested in a catheter-associated IE model (Cheung et al., 1994; Seidl et al., 2011). Yet, with a deficient Agr system, IE1789 virulence is severely diminished. Heart valve infection fails to promote dissemination and acute kidney injury, resulting in most of the rabbits surviving the experimental period in spite of the presence of large vegetations. Without the RNAIII-regulated exoproteins, *S. aureus* native valve IE resembles that of the oral pathogen *Streptococcus sanguinis*, characterized by subacute, chronic thrombosis with low systemic toxicity and low lethality (Martini et al., 2020).

In stark contrast, BA0211 IE leads to metastatic infection, the most severe kidney injury, and lethality in all experimental rabbits. *RNAIII* expression occurs at a slower rate resulting in overall low expression with concomitant high *rot* and *sarS* expression. In BA0211, *sarA* expression is not only low but also uninduced throughout growth, partially explaining the lower RNAIII levels observed in this strain and decreased cytotoxicity towards rabbit erythrocytes. The expression patterns of the global regulators in this strain suggest that large aortic valve vegetations accompanied by severe systemic toxicity arise from strains that express colonization factors at high level but have a slower transition towards expression of the secreted virulence factors.

Overall, we cannot exclude the possibility that other regulatory elements are responsible for the clinical outcomes of the *S. aureus* CC5 isolates tested in our model. We also acknowledge that expression can vary between *in vitro* and *in vivo* settings. We are poised to address this in future studies. In conclusion, this study highlights the requirement of a more measured expression of *RNAIII* and *sarA* as an intrinsic phenotypic characteristic of strains proficient in development of IE with severe complications. Simultaneous high expression of *RNAIII, sarA*, *sigB*, and *mgrA* leads to severe systemic toxicity but is the one phenotype that fails to promote vegetation formation in the native valve model. Thus, *RNAIII* and *sarA* expression that provides for rheostat control of colonization and virulence genes, rather than an on and off switch, promote both vegetation formation and lethal sepsis.

## Author Contributions

Conceptualization, K.J.K. and W.S.-P.; Methodology, K.J.K., K.N.G.-C., and W.S.-P.; Formal Analysis, K.J.K. and W.S.-P.; Investigation, K.J.K., J.M.S., K.K, M.B., and K.N.G.-C., and W.S.-P.; Writing – original draft, K.J.K. and W.S.-P.; Writing – review & editing, K.J.K., J.M.S., K.K, M.B., and K.N.G.-C., and W.S.-P.; Visualization, K.J.K. and W.S.-P; Supervision, W.S.-P.; Funding Acquisition, W.S.-P.

## Acknowledgments

We thank Katherine N. Gibson-Corley, DVM, Ph.D., Dipl. ACVP for providing gross pathology expertise, and François Vandenesch, M.D., Ph.D. for providing the *S. aureus* CC5 isolates.

