## Supplemental Materials for "Vegetation formation in *Staphylococcus aureus* endocarditis inversely correlates with *RNAIII* and *sarA* expression in invasive clonal complex 5 (CC5) isolates"

**Supplementary Material**

**Supplementary Figures and Tables**

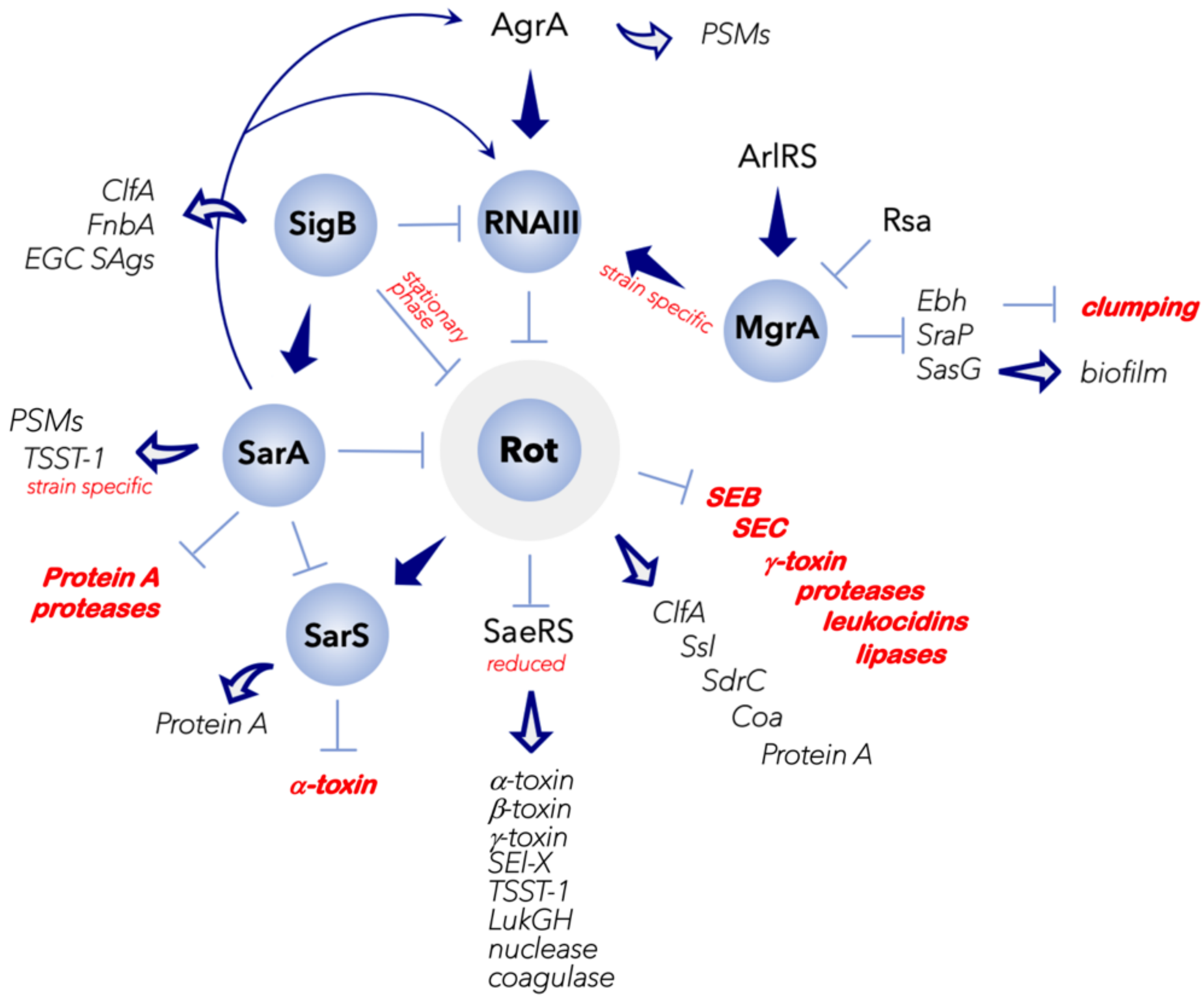


**Figure S1. Current understanding of *S. aureus* global regulator control of gene expression.** The Agr quorum-sensing system differentially controls expression of *S. aureus* surface proteins [e.g. microbial surface components recognizing adhesive matrix molecules "(MSCRAMMs) and protein A] and the secreted toxins and enzymes (e.g. hemolysins, superantigens, proteases, nucleases, lipases) (Jenul and Horswill, 2019). RNAIII is the effector molecule of the Agr system and its expression is regulated by AgrA. When the Agr system is inactive, *rot* (for *repressor of toxins*) is expressed. Rot induces SarS and surface proteins and reduces SaeRS and secreted toxins and enzymes. When the Agr system is active, *RNAIII* is expressed and Rot is translationally suppressed. Inhibition of Rot allows for expression of the secreted toxins and enzymes and concomitantly reduces expression of SarS and surface proteins (Mcnamara et al., 2000; Saïd-Salim et al., 2003; Geisinger et al., 2006). SarA increases expression of the *agr* system (Heinrichs et al., 1996; Rechtin et al., 1999) and genes encoding for the phenol-soluble modulins (PSMs) (Morrison et al., 2012) and represses expression of *sarS* and *rot* (Cheung et al., 2001; Hsieh et al., 2008). SarS acts opposite to SarA by inducing expression of *spa* (Protein A) and repressing expression of several toxin genes such as *hla* (α-toxin gene) (Tegmark et al., 2000). The alternative sigma factor B (SigB) directly inhibits expression of the *agr* operon (Bischoff et al., 2001) and of *rot*

during growth in stationary phase (Hsieh et al., 2008), but induces expression of *sarA*, adhesin genes and the enterotoxin gene cluster (*egc*) (Bischoff et al., 2004; Entenza et al., 2005; Kusch et al., 2011). MgrA regulatory effects mirror those of the *agr* system, although the specific targets can be distinct (Luong et al., 2006). MgrA upregulates production of secreted proteins (e.g. leukotoxins, Spl proteases, enzymes) and downregulates production of surface-associated proteins (e.g. large surface protein Ebh, SraP, SasG) (Luong et al., 2006; Crosby et al., 2016).

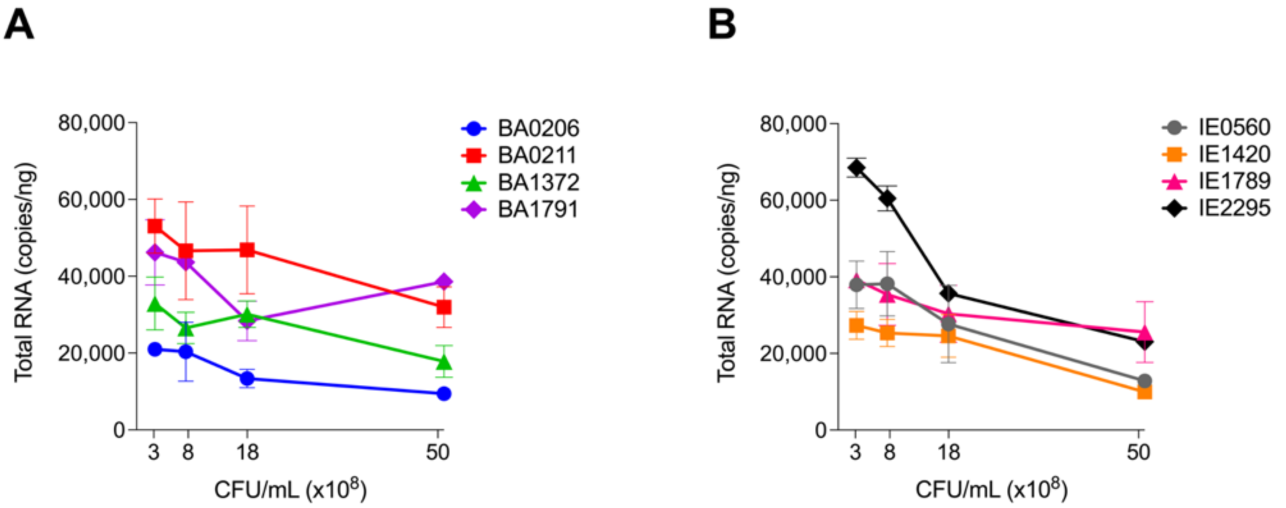


**Figure S2. CC5 clonal group differentially express *gyrB*.** (**A, B**) Quantitation of *gyrB* mRNA in *S. aureus* CC5 during growth in TH broth by RT-qPCR standard curve quantitation method. Data is represented by the mean ± SEM of three biological replicates. Error bars (standard deviation) not shown are smaller than symbol. Two-way ANOVA with Holm-Šídák's multiple comparisons test.

**Table S1. Kidney gross pathology grading scale**

| **Kidney Pathology** | | |
|---|---|---|
| Color: red/hemorrhagic (+/-) white/grey/necrotic (+/-) mottled red/grey (+/-) | 0 | No lesions |
| | 1 | Rare, up to 4-5 small (<4mm) multifocal lesions (infarcts) on surface |
| | 2 | Numerous larger (>5mm) multifocal lesions (infarcts) on surface |
| | 3 | Locally extensive to coalescing to diffuse lesions (infarcts) on surface |

**Table S2. RT-qPCR Primers**

| Primer | Primer Sequence |
|---|---|
| gyrB_RTqPCRFwd<br>gyrB_RTqPCRRev | CGTCCAGCTGTCGAAGTTATT<br>CTGATGAACCAACACCATGTAAAC |
| RNAIII_RTqPCR_For<br>RNAIII_RTqPCR_Rev | GCACTGAGTCCAAGGAAACT<br>AGCCATCCCAACTTAATAACCA |
| rot_RTqPCR_For<br>rot_RTqPCR_Rev | GCATTGCTGTTGCTCTACTTG<br>CGTCCTGTTGACGATGAAAGA |
| mgrA_RTqPCR_For<br>mgrA_RTqPCR_Rev | TCGGAACGTTCACGCTTAAT<br>TCTCCTGTAAACGTCAAGAAAGT |
| SigB_RTqPCR_For<br>SigB_RTqPCR_Rev | TCGCGAACGAGAAATCATACA<br>CCGTTCTCTGAAGTCGTGATAC |
| SarA_RTqPCR_For<br>SarA_RTqPCR_Rev | TAGCTTTGAAGAATTCGCTGTATTG<br>GCTTTAACAACTTGTGGTTGTTTG |
| SarS_RTqPCR_For<br>SarS_RTqPCR_Rev | CGCTCAACTGAAGATGAAAGAAA<br>TGAGCTAATAATTGTTCAGCATGG |
| SEG RT-qPCR For<br>SEG RT-qPCR Rev | ACTGTACAGGTAACAATCGACAATAG<br>TGCAGAACCATCAAACTCGTATAG |
| SEl-I RT-qPCR For<br>SEl-I RT-qPCR Rev | GGGCCACTTTATCAGGACAATAC<br>ACATCAATTTCTTGAGCTGTGACTA |
| SElM_N315_RTqPCR_For<br>SElM_N315_RTqPCR_Rev | GGTGGAGTTACATTAGCAGGTG<br>TGATGTTCTCCATTAACCCAAAGA |
| SEl-N RT-qPCR_ For<br>SEl-N RT-qPCR_ Rev | GGACTGTATTATGGAAATAAATGTGTAGGC<br>ACCTTCTTGTTGGATACCATCTT |
| SEO_N315_RTqPCR_For<br>SEO_N315_RTqPCR_Rev | TTTAGCTCATCAGCGATTTCTAAAG<br>CCACCATATGTACAGGCAGTAT |
| SEl-U_RTqPCRFwd<br>SEl-U_RTqPCRRev | GTGTTAAGTCTTGCAGCTTACTATT<br>CCTCATATTATCCATTAGACCAGTGA |
| SEl-X_RTqPCRFwd<br>SEl-X_RTqPCRRev | TCTATCGCTAGGTATCATCTATGGG<br>GGAATTGTTTATCTTGTACACTTGGG |

### Table S3. qPCR assay performance

| Target Amplicon | Amplicon Length (bp) | Melt Curve Tm (SD) | PCR Efficiency (%) | $R^2$ Calibration Curve | Linear Dynamic Range (Copies) | Intraassay Variance (SD of $C_q$) |
|---|---|---|---|---|---|---|
| gyrB | 103 | 80.56 (0.28) | 97.73 | 0.999 | 100-1,000,000 | 0.06 |
| RNAIII | 81 | 76.69 (0.15) | 103.5 | 0.998 | 100-10,000,000 | 0.08 |
| rot | 118 | 77.99 (0.14) | 101.1 | 0.996 | 100-1,000,000 | 0.06 |
| mgrA | 113 | 78.22 (0.14) | 99.02 | 0.999 | 100-1,000,000 | 0.09 |
| sigB | 110 | 79.34 (0.15) | 99.84 | 0.999 | 100-1,000,000 | 0.12 |
| sarA | 114 | 75.22 (0.15) | 97.65 | 0.999 | 100-1,000,000 | 0.08 |
| sarS | 80 | 77.32 (0.14) | 99.33 | 0.999 | 100-1,000,000 | 0.09 |
| seg | 147 | 77.56 (0.17) | 98.92 | 0.998 | 10-1,000,000 | 0.04 |
| sei | 139 | 77.27 (0.17) | 100.5 | 0.999 | 10-1,000,000 | 0.06 |
| sem | 80 | 74.72 (0.15) | 101.3 | 0.998 | 10-1,000,000 | 0.07 |
| sen | 150 | 77.37 (0.21) | 102.6 | 0.998 | 10-1,000,000 | 0.06 |
| seo | 119 | 78.08 (0.12) | 98.41 | 0.998 | 10-1,000,000 | 0.13 |
| se*l*u | 101 | 76.83 (0.05) | 96.4 | 0.998 | 100-1,000,000 | 0.10 |
| se*l*x | 98 | 78.51 (0.14) | 101.3 | 1.000 | 100-1,000,000 | 0.09 |
